# Deciphering the Functional Roles of Individual Cancer Alleles Across Comprehensive Cancer Genomic Studies

**DOI:** 10.1101/2023.11.14.567106

**Authors:** Jiayan (Yoshii) Ma, Stephanie Ting, Bartholomew Tam, Timothy Pham, Michael Reich, Jill Mesirov, Pablo Tamayo, William Kim

**Affiliations:** Center for Novel Therapeutics, UC San Diego, La Jolla, CA, 92037; Moores Cancer Center, UC San Diego, La Jolla, CA, 92093; School of Medicine, UC San Diego, La Jolla, CA, 92093; Division of Genomics and Precision Medicine, UC San Diego, La Jolla, CA, 92093

## Abstract

Cancer genome data has been growing in both size and complexity, primarily driven by advances in next-generation sequencing technologies, such as Pan-cancer data from TCGA, ICGC, and single-cell sequencing. Yet, discerning the functional role of individual genomic lesions remains a substantial challenge due to the complexity and scale of the data. Previously, we introduced REVEALER, which identifies groups of genomic alterations that significantly associate with target functional profiles or phenotypes, such as pathway activation, gene dependency, or drug response. In this paper, we present a new mathematical formulation of the algorithm. This version (REVEALER 2.0) is considerably more powerful than the original, allowing for rapid processing and analysis of much larger datasets and facilitating higher-resolution discoveries at the level of individual alleles. REVEALER 2.0 employs the Conditional Information Coefficient (CIC) to pinpoint features that are either complementary or mutually exclusive but still correlate with the target functional profile. The aggregation of these features provides a better explanation for the target functional profile than any single alteration on its own. This is indicative of scenarios where several activating genomic lesions can initiate or stimulate a key pathway or process. We replaced the initial three-dimensional kernel estimation with multiple precomputed one-dimensional kernel estimations, resulting in an approximate 150x increase in speed and efficiency. This improvement, combined with its efficient execution, makes REVEALER 2.0 suitable for much larger datasets and a more extensive range of genomic challenges.

## Introduction

Over the last decade, systematic efforts to sequence large numbers of tumors have provided a rich catalog of the most common genetic alterations that drive cancer initiation and maintenance.^1,2^ However, the task of identifying the functional role of these lesions is fraught with difficulties. Genomic instability, an increased rate of mutations occurring throughout the genome, adds a layer of complexity to the study of cancer genetics as it results in an increased number of genomic alterations. Included in these are alterations with low penetrance. These events can be particularly challenging to characterize due to their low prevalence, to determine exactly how these low penetrance events contribute to the overall cancer phenotypes, and to distinguish between the more common mutations with functional significance. To complicate matters, the role of the same genetic alteration can be different according to the specific biological context or cellular state in which it takes place. This contextuality compounds the already challenging task of functionally annotating and interpreting the genetic landscape of cancers and identifying key events critical for the development and progression of the disease. ^3–5^

We previously introduced REVEALER,^6^ (*Repeated Evaluation of VariablEs conditionAL Entropy and Redundancy*) to address some of these challenges. REVEALER is an algorithm that identifies groups of genomic alterations that have significant association with a biological phenotype of interest representing, for example, pathway activation, gene dependency, or drug response. REVEALER makes use of powerful Information-Theoretic measures of association, the *mutual information* and the *conditional mutual information*,^*7*^ to identify combinations of these alterations that explain the biological phenotype better than any individual alteration considered in isolation. REVEALER can identify features that are complementary or mutually exclusive, yet still correlate with a target functional profile. In this scenario multiple activating genomic lesions can induce or activate a driver pathway or process, but they rarely appear together since a single event is sufficient to activate or induce the functional target. Mutual exclusivity, while not without exceptions, is useful in practice. To demonstrate the value of the approach in *Kim et al 2016*^*6*^, we showed that REVEALER can be used to explore the mutational landscape associated with an specific oncogenic pathway activation, drug response or any other functional readouts, and identify relevant genomic features with higher sensitivity and specificity compared to other feature selection methods.^6^ We explored the mutational and CNA landscape of complementary genomic alterations associated with the transcriptional activation of β-catenin, NRF2, and with MEK-inhibitor sensitivity, and KRAS dependency. In all these cases REVEALER successfully identified both known and novel associations, demonstrating the power of the approach that combines functional profiles with cancer datasets containing extensive characterization of genomic alterations.

Since its publication, REVEALER has been applied to a variety of problems; for example, we have used it to identify FAT1 truncating mutations as the top genomic abnormality complementary of YAP1 gene amplification significantly associated with YAP1 activity.^8^ It has also been used to systematically find the set of genomic alterations, copy number and mutations, that are most likely responsible for the dysregulation of Immune-Associated (IA) genes in hepatocellular carcinoma (HCC).^9^ In another application, *REVEALER* was used to correlate all genomic alterations present in bladder cancer samples to the expression of selected dysregulated miRNAs^10^. Despite its successful application to problems at the forefront of cancer research, such as the ones listed above, REVEALER’s original implementation, using R-language libraries for 3D kernel density and bandwidth estimation, has relatively large execution times limiting its applicability to large datasets such as TCGA.^11^ In this pape*r we introduce a completely new mathematical formulation of the algorithm and expand its use modalities (REVEALER 2*.*0)*. We replaced the 3D kernel estimation of the original implementation with several one-dimensional kernel estimations, using precomputed Gaussian distribution kernels resulting in a *150-fold* increase of performance. In addition, we introduced novel pre-processing options that enable its application to a wider repertoire of applications, including the analysis of individual alleles. The new and expanded formulation of the algorithm and its efficient implementation enable the use of REVEALER 2.0 in much larger datasets and in a wider class of genomic problems, including the ability to rapidly run the analysis with multiple different parameters, both at the level of individual alleles and copy number alterations. In the next section we will review the basics of the algorithm, its new mathematical formulation, implementation, examples of use, and performance improvements and benchmarks.

## Results

### Overview of the Algorithm

There are three inputs to REVEALER 2.0: 1) a “target” profile (*t*), typically a readout representing a functional phenotype such as a gene expression, pathway activation, gene-dependency or drug response profile across a group of samples; 2) a dataset (*F= {f*_*i*_*}*) containing a comprehensive collection of binary genomic “features” *f*_*i*_, for the same samples where the target profile is defined, and; 3) an *original* “*seed*” (*s*), an optional binary feature to initialize the search, which is augmented at each REVEALER 2.0 interaction and becomes a *new seed (s*_*j*_). The seed is typically a feature that is known to be an activating “cause” of the target profile such as a mutation of a driver oncogene. REVEALER 2.0 makes use of two powerful Information-Theoretic metrics, the *Information Coefficient* (IC) used to quantify the degree of association between the *original* or *new seed* and the *target*, and the *Conditional Information Coefficient* (CIC) to quantify the degree of association between a *feature* and the *target* conditional to the value of the *original* or the *new seed*.

**Fig. 1**. shows a summary of the algorithm at the top level. REVEALER 2.0 performs a number of iterations where it tries to find the *feature* most associated (correlated) with the *target* that is also complementary to the *seed*. In the first iteration, it computes the CIC between the *target* and all the input *features* conditional to the *original seed* (Fig.1A). Once it finds the *feature* with the highest CIC (*e*.*g. feature f*_*1,1*_ in 1A), it combines it with the seed (binary addition) to generate a *new seed* (Fig.1B) and start a new iteration (Fig.1C). This process is repeated (Fig.1D) until the IC between the target and the *new seed stops* increasing (Fig.1E). The final result consists of the top matching features from each iteration (Fig.1 F) and the final IC and corresponding *seed* that combines all the top matching features. The top matching features together explain the target’s activation better than any of them individually (Fig. 1G).

**Fig. 1.**
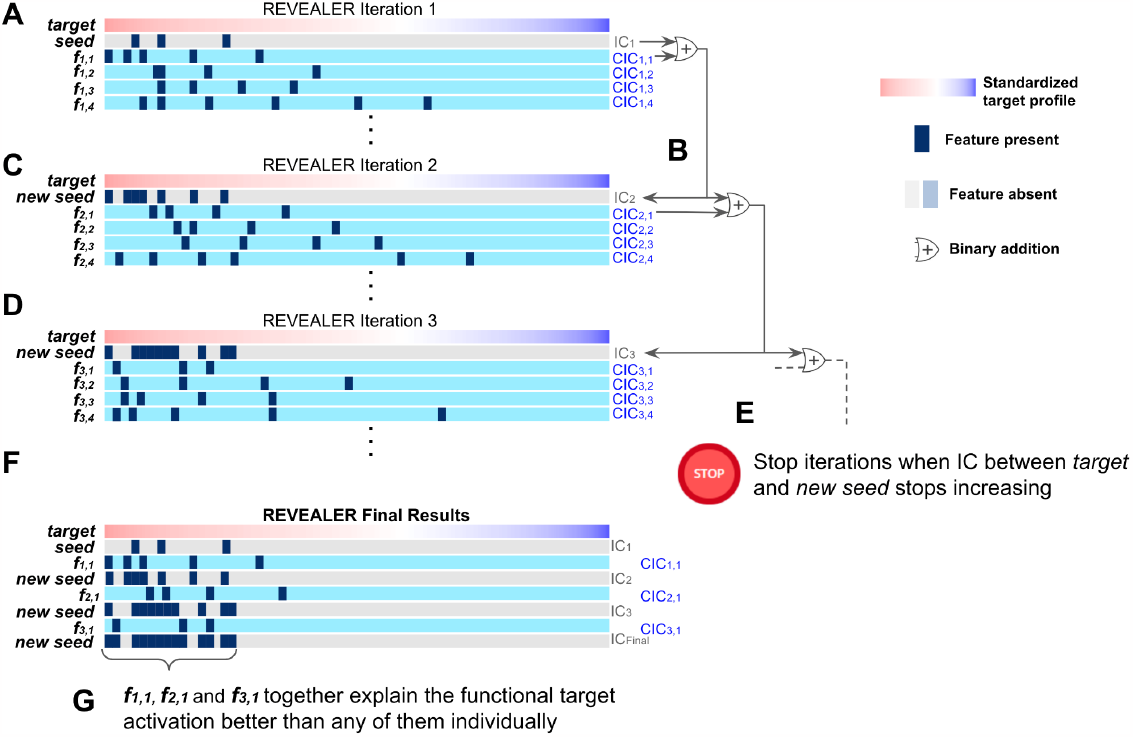
Schematic diagram of the REVEALER algorithm’s iterative scheme and final results.

In REVEALER 2.0, the outer iterations over features are implemented as in the original version, however the estimation of the *mutual* and *conditional mutual information* are optimized so that those quantities can be efficiently computed using a discrete grid where the relevant density probability distributions can be approximated with precomputed kernels representing Gaussian distributions.

### New Formulation of the REVEALER Computation

Let’s start with definition of the *mutual information*^*7*^ that measures the degree of association between a *target* (*t*) and a *feature* (*s*),

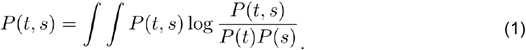

The *features* in our case are binary variables, e.g. the mutation or amplification/deletion status of genes, and the *target*, a *functional profile* of interest, is continuous. The target can be discretized so that the two variables are discrete and the computation can be executed using a discrete grid of dimensions *n*_*t*_ x 2, where there are only *n*_*t*_ distinct values for the target (see top of Fig. 2). Taking this approximation into account the definition of the mutual information for discrete variables is, ^7^

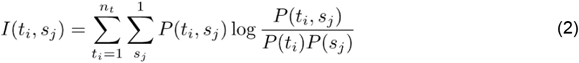

where the target takes one of *n*_*t*_ values in each sample and the binary variable *s*_*j*_ is either the original seed (*s*_*0*_), or the new seed (*s*_*j*_) after each iteration of the algorithm, and takes only two values (0, 1). REVEALER 2.0 makes use of *The Information Coefficient (IC)*, a rescaling of the *mutual information* that takes values between 1 (perfect match) and −1 (perfect anti-match),^6,12^

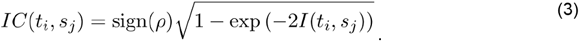

**Fig. 2.**
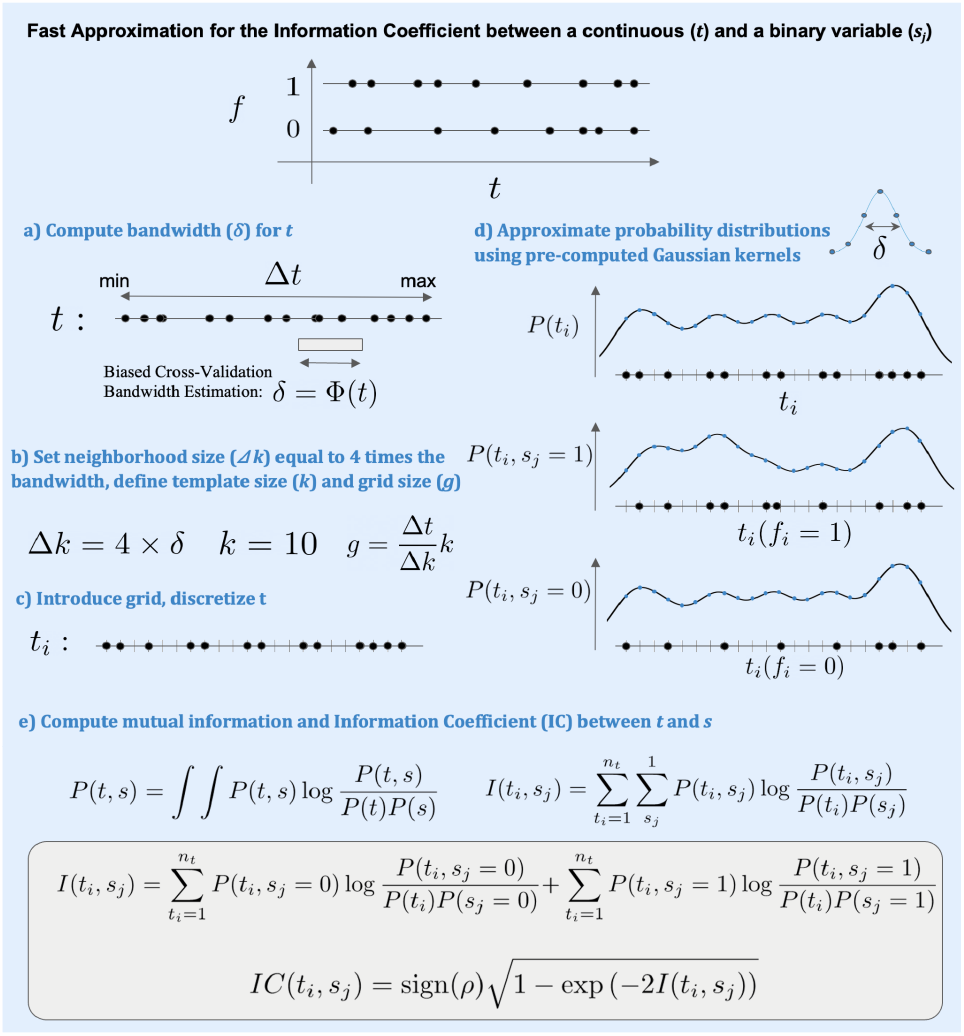
Procedure to implement a fast approximation for the *Information Coefficient (IC)* between the *target* (*t*) and a binary variable (seed, *s*_*j*_).

Notice that in this expression the sign of the standard correlation coefficient *ρ(t, f*_*i*_), between the target and the binary variable, is used to provide *directionality*. In REVEALER 2.0 we clearly require directionality in order to distinguish between positive and negative association with the target.

The procedure to approximate the *Information Coefficient (IC)* consists of the following steps as shown in **Fig. 2**: First, a) compute the bandwidth of the *target* variable (Fig. 2A) using a fast Cython implementation of a biased-cross validation bandwidth estimation algorithm.^13^ This bandwidth will be used to parameterize the kernels; b) compute the neighborhood size (Fig. 2B); c) introduce a discrete grid and discretize the *target* (Fig. 2C); d) approximate the probability distribution on the grid using pre-computed Gaussian kernels (Fig. 2D). For each data point a kernel is placed at the corresponding grid point and the kernel densities are added to their corresponding grid point and every grid point in the neighborhood. Finally, e) Compute the *Information Coefficient (IC)* (Fig. 2E).

The described procedure allows for a fast computation of the *IC* and the same strategy will be used to compute somewhat more complex *Conditional Information Coefficient (CIC)* which is the workhorse of REVEALER 2.0 ‘s iterations. The *CIC* is closely related to conditional mutual information,^7^

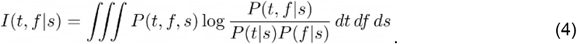

Which can be also defined for discrete variables as follows,^7^

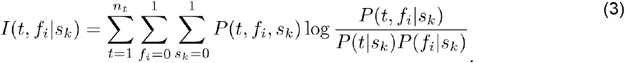

In this case the computation can be implemented using a three-dimensional grid of dimensions *n*_*t*_ x 2 x 2 (top of Fig. 3). The *Conditional Information Coefficient (CIC)* is defined in an analogous way to the *Information Coefficient*.

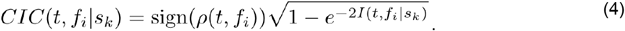

**Fig. 3.**
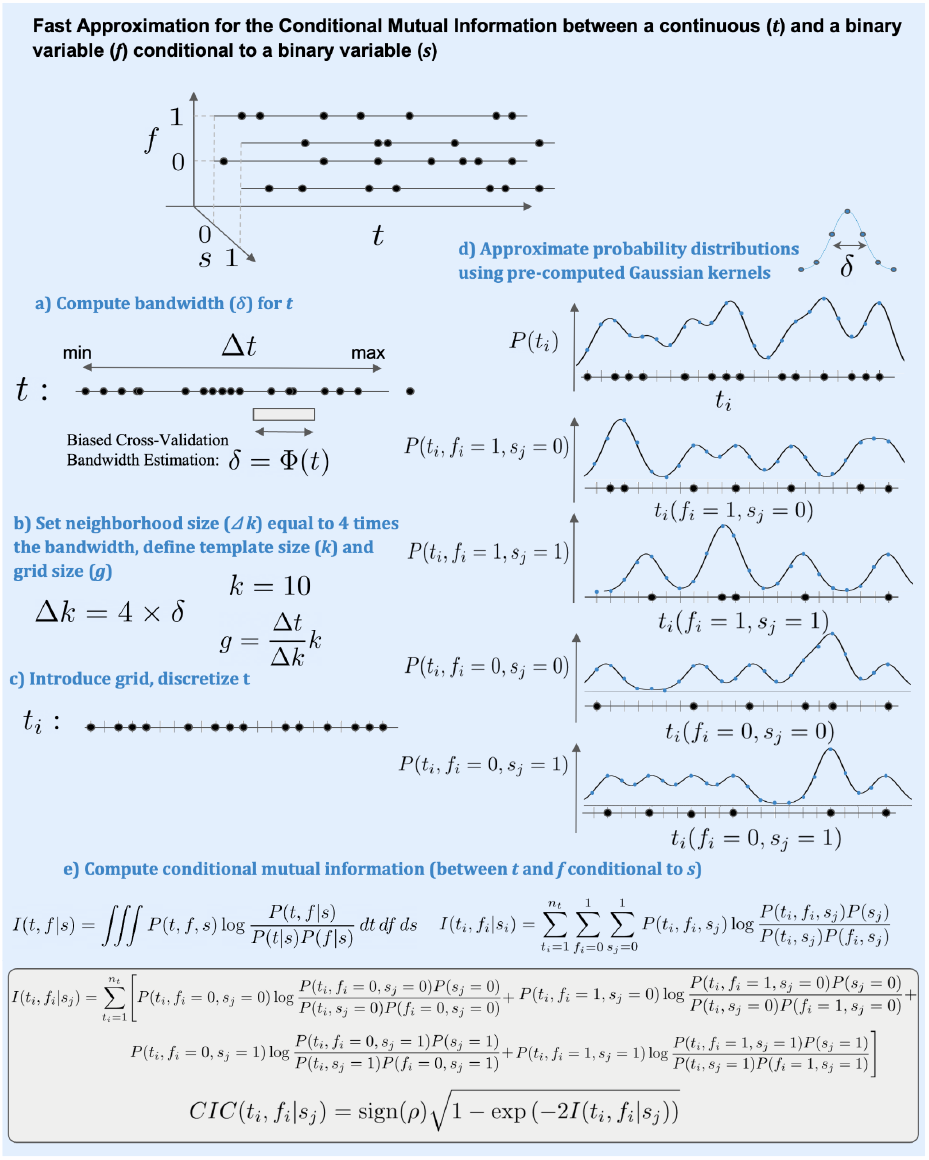
Procedure to implement a fast approximation for the *Conditional Information Coefficient (CIC)* between the *target* (*t*) and a binary variable (feature, *f*_*i*_) conditional to a second binary variable (seed, *s*_*j*_).

The directionality, as in the case of the *IC*, is determined using the standard correlation coefficient *ρ(t, f*_*i*_), between the target (*t*) and the feature binary variable(*f*_*j*_). In a similar way to the approximation for the Information Coefficient (IC), the procedure to approximate the *Conditional Information Coefficient (CIC)* consists of the following steps: a) compute bandwidth for the *target* variable (Fig. 3a); b) Compute neighborhood size (Fig. 3b); c) introduce a discrete grid and discretize the *target* (Fig. 3c); d) approximate the probability distribution on the grid using pre-computed Gaussian kernels (Fig. 3d); e) Compute the *Conditional Information Coefficient (CIC)* (Fig. 3e). Notice how the final computation of the *IC* and *CIC*, denoted in gray boxes at the bottom of Figs. 2 and 3, can be performed very efficiently under the proposed approach. In the next section we will provide application examples of REVEALER 2.0 and we will describe some of the benchmarks we used to test the correctness of the implementation and to assess its performance and properties.

### Validation benchmarks and performance measurements

Synthetic data corresponding to different known distributions was generated to test the validity or REVEALER 2.0 in its approximations and implementation, and also to help determine default values for parameters such as the overall multiplicative kernel bandwidth.

1. **Bandwidth estimation**. We implemented the bandwidth calculation using a fast Cython implementation of a biased-cross validation algorithm^13^ which is the same as used in the original implementation of REVEALER (the bcv function in *R’s* package MASS^14^). We compared values of bandwidth estimation for continuous target with a variety of distributions (uniform, normal, exponential, binomial and poisson) and managed to reproduce the values produced by *R’s* bcv bandwidth estimation used in the original REVEALER implementation (Fig.4A).
2. **Exact analytical integration of IC for a piecemeal uniform distribution: Fig. 4B**. shows how REVEALER 2.0 agrees with the IC results obtained by analytically integrating the relevant probability distributions and computing the entropy.
3. **Numerical Integration ofGaussian and Sigmoidal conditional distributions to estimate the IC and CIC**. In this case we define a normally distributed target while the features are binary variables with sigmoidal conditional probabilities. In **Fig. 4C-D** we can see that REVEALER 2.0’s estimates for the IC and CIC from these types of synthetic data agree well with the computations of those quantities using numerical integration of the entropy.
4. **REVEALER 2.0 execution times**. The more efficient formulation and implementation of REVEALER 2.0 takes advantage of using discretization and precomputed kernels and also benefits from using Cython compiled code to execute the most expensive procedures such as the computation of the kernel bandwidth. In **Fig. 5**, we show typical running times for REVEALER 2.0 and the original implementation in *Kim et al*.*2016*.^*6*^ The different feature size datasets shown in the figure were obtained by subsetting one of the TCGA mutation files (102,326 rows × 8,671 columns) into subsets of 250, 2,500, 10,000 and 100,000 features and keepings all the columns. REVEALER 2.0’s execution time for the entire TCGA dataset (with over 100,000 features) is approximately 45 minutes. We estimate that it will take 137 hours (app 6 days) if run on the same machine using the original implementation,^6^ making REVEALER 2.0’s implementation about 185 times faster.

**Fig. 4.**
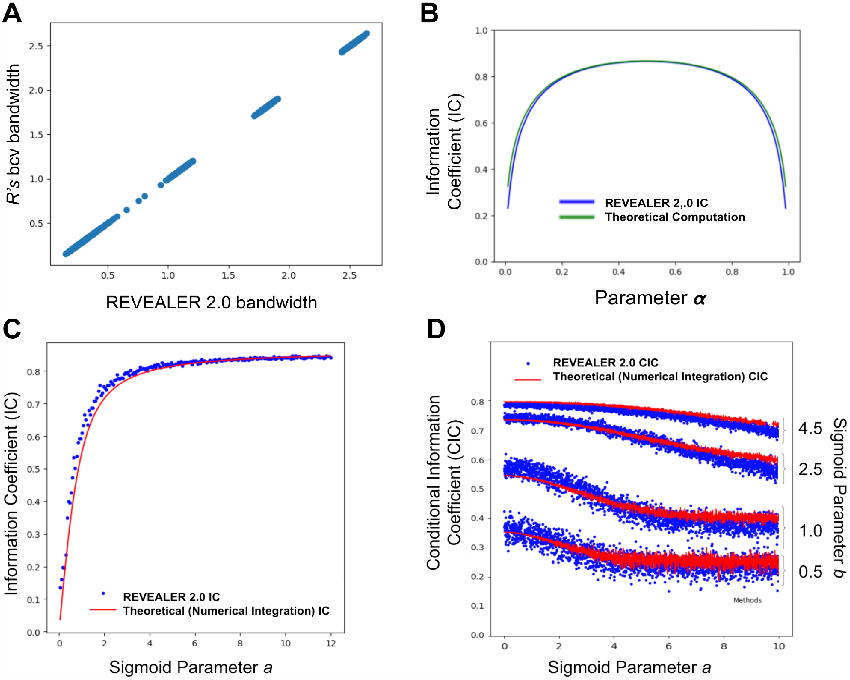
Benchmarks to validate the correctness of the algorithm’s approximation. A) comparison between *R’s* bcv bandwidth estimation^14^ and REVEALER 2.0. B). Comparison of REVEALER 2.0’s IC estimate vs. theoretical computation using a uniform piecemeal distribution. C-D) IC and CIC comparison of REVEALER 2.0 vs. theoretical/numerical Integration for a Gaussian distributed target and sigmoid distributed conditional probabilities.

**Fig. 5.**
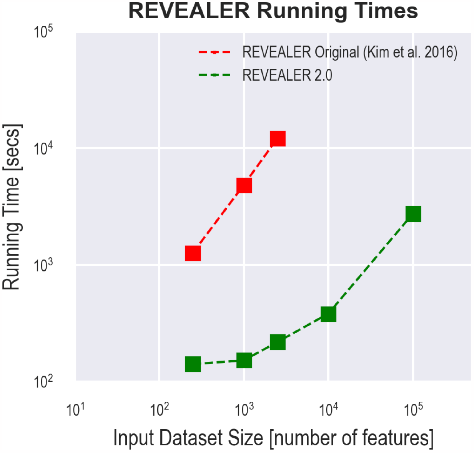
Running times for the original and this paper implementations of REVEALER.

### Application Examples

To highlight the new capabilities of REVEALER 2.0, we focused the analysis on individual alleles (vs. copy number alterations) as examples.

### MEK inhibitor sensitivities

Dysregulation of the MAPK signaling pathway is known to contribute to one of the major hallmarks of cancer, and this has led to the development and clinical testing of several MAPK inhibitors.^15^ While there are a number of well-known oncogenic mutations in upstream kinases that are responsible for activating this pathway, the ability to pinpoint specific mutations that can be used to predict sensitivities to clinical inhibitors of the pathway has immediate therapeutic implications. We and others have previously demonstrated that a gene expression signature can be used to predict the response to MAPK inhibitors.^16,17^ We therefore used this signature score as a functional profile to assess whether we can use REVEALER 2.0 across a pan-cancer TCGA dataset (10,956 samples, 32 tissues) to discover specific alleles that may be associated with MAPK inhibitor sensitivities. Indeed, upon running the analysis, the top scoring alleles are many known activating alleles in BRAF, KRAS and NRAS^18–20^ (Fig. 6). We also applied this analysis across the *Cancer Cell Line Encyclopedia* (CCLE^21^) and found that KRAS_p.G12D, KRAS_p.G12V, NRAS_p.Q61K alleles were also among those scoring as the top hits from this analysis (Fig. S1). To note, while many alleles which are less frequent (n < 3) and did not score highly, the analysis was able to discern alleles that are variants of known hotspots (e.g., BRAF_V600E), suggesting that the analysis can help identify low frequency alleles with functional relevance. We also found that there were number outlier samples even within the top scoring alleles, suggesting that in the absence of functional readouts, relying on individual alleles entirely to predict response to drugs may be insufficient.

**Fig. 6.**
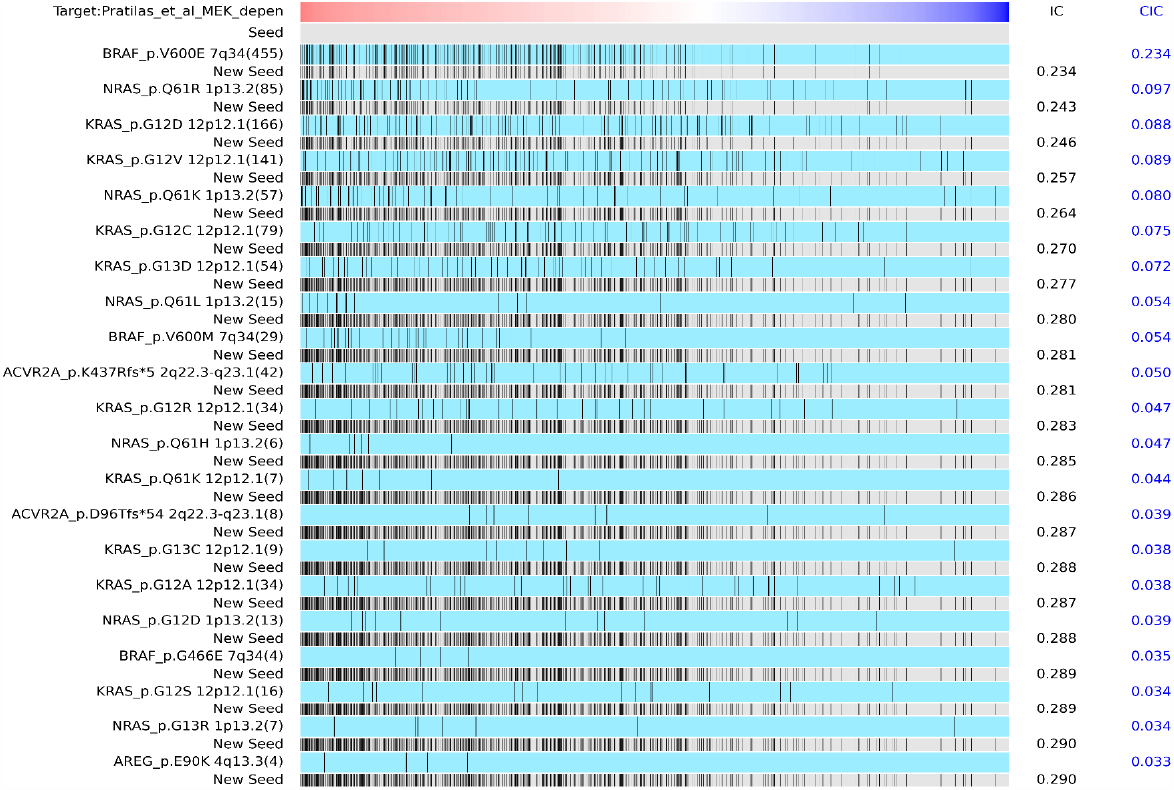
Individual alleles identified to match MEK inhibitor sensitivity profiles across TCGA pan-cancer dataset (10,956 samples, 32 tissues).

We also observed two novel events, including hotspot mutations in AREG and ACVR2A, which were significantly associated with MAPKi sensitivities in CCLE and TCGA datasets, respectively (Fig. 6, Fig. S1). AREG is a member of the epidermal growth factor (EGF) and has been shown to activate the MAPK directly via engagement with the EGF receptor^22^ (Fig. 6). We also found frameshift mutation in the gene ACVR2A (K437Rfs) (Fig. 6). ACVR2A is an activin receptor 2A, a member of the TGF-b superfamily^23^. While there are no immediate reports of ACVR2A direct engagement of the MAPK pathway, there are reports which suggest that it may engage the MAPK pathway indirectly via SMAD family of proteins^24^. Taken together, these results highlight new opportunities to further identify additional genetic alterations that may shed new biological insights underlying the MAPK pathway activation and its inhibition.

### β-catenin/WNT pathway

Activation of β-catenin/WNT pathway *via* loss of APC and/or activating mutations in β-catenin occurs at high frequencies across many cancer types, including Colorectal, Hepatocellular Carcinoma, and Endometrial Cancer.^25^ Identifying which mutant alleles are functional and associated with activation of the pathway is a challenge because there are a large number of alleles observed in the cancer genome. We applied REVEALER 2.0 to identify β-catenin and APC mutant alleles which can activate WNT pathway using β-catenin/WNT hallmark signature score across pan-cancer TCGA.^6,26,27^ In TCGA, there are in total 169 β-catenin and 587 APC alleles. Out of these, we identified 306 alleles to be highly commentary among each other and associate with β-catenin/WNT pathway activation (Fig. 7). These included mutations in the serine/threonine residues of β-catenin at positions 33, 37, 41, and 45, and 1450, 213, and 216 residues in APC. Out of all 587 alleles in APC and 169 alleles in β-catenin observed in TCGA, 38 and 23 were also found in CCLE, respectively (Fig. S2-S3, TABLE S1). Among these, 28 APC and 15 β-catenin alleles were also identified as top hits in CCLE, suggesting that many of these lesions may be functionally relevant. As with the MEK inhibitor example (Fig. 6), there are several alleles which only occur once in each dataset and assessing functional significance of these alleles require follow-up studies. The results of the REVEALER 2.0 can guide prioritization of these alleles.

**Figure 7.**
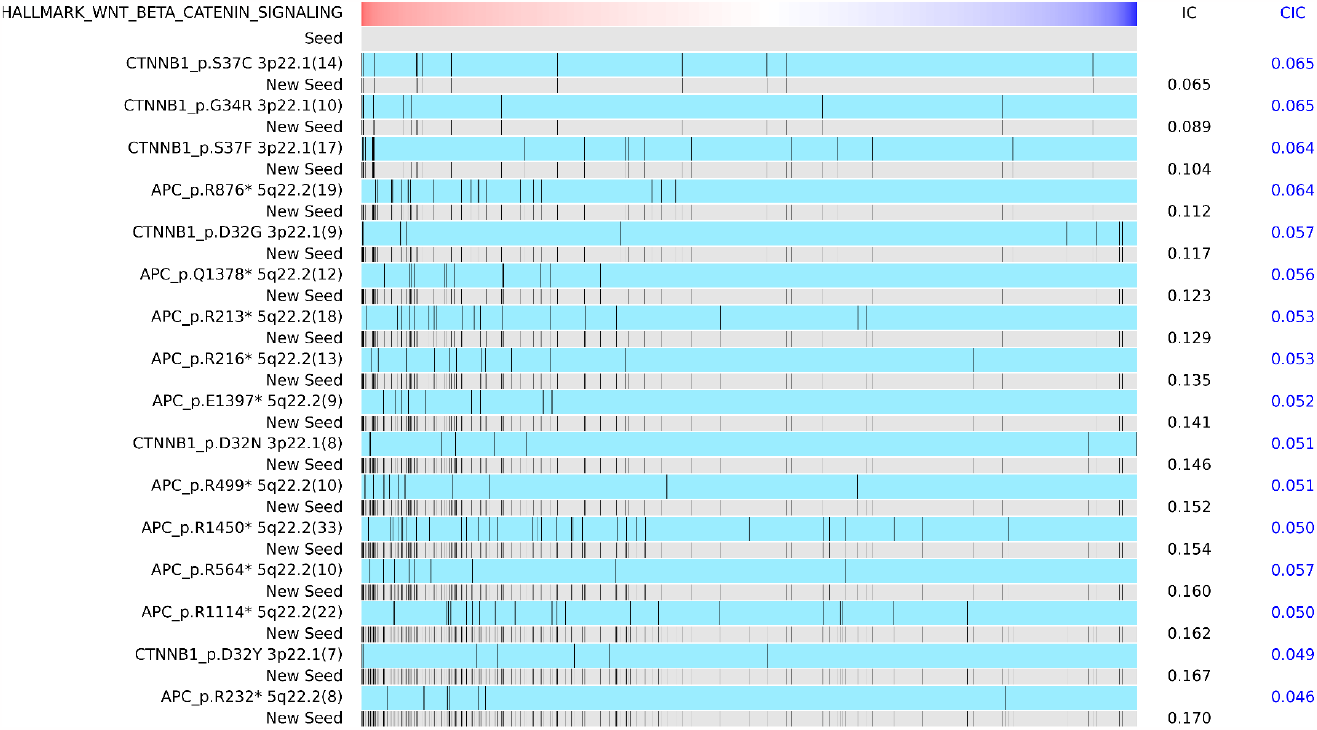
Individual alleles of β-catenin and APC identified to match β-catenin/WNT pathway activation profiles across TCGA pan-cancer dataset.

We then used the β-catenin and APC alleles, which are found in both TCGA and CCLE datasets and are positive correlated, as seeds, and ran the REVEALER 2.0 to identify other mutations which may be complementary to β-catenin and APC (Fig.7B). Among the top hits were one of the β-catenin allele (G34R). This is notable as this allele was not included in the original seed because it was not represented in the CCLE. This result shows that REVEALER is sensitive in identifying relevant alleles. Also among the top hits were GTF2I, TBP, GNAQ, GNA11, and TMPRSS13 (Fig. 8). GTF2I and TBP are general transcription factors which have been implicated^28^ in the β-catenin/WNT pathway, or shown to biochemically interact with β-catenin,^29–31^ respectively. GNAQ and GNA11 are G-protein coupled receptors which are predominantly found in Uveal melanoma.^32^ Recent studies suggest that mutation GNAQ is upstream of the YAP1,^33^ which has also been shown to be a transcriptional co-activator of β -catenin.^34^ TMPRSS13 is a cell surface protease whose mutations have been found in colorectal cancer, UVEAL and cutaneous melanoma.^32^ However, its precise roles are unknown. It is plausible to postulate, based on the REVEALER 2.0 results, that these genes, and the corresponding mutations may be involved in the β-catenin/WNT pathway activation.

**Figure 8.**
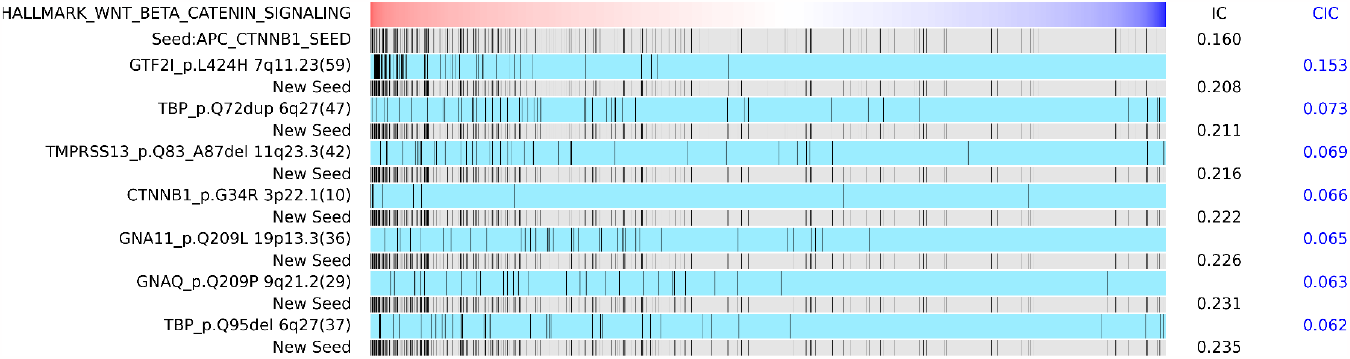
Individual alleles identified to match β-catenin/WNT pathway activation profiles across TCGA pan-cancer dataset.

In summary, these examples illustrate that REVEALER 2.0 is capable of discoveries which may have been difficult to implement in the earlier version. Specific modalities of use cases described here not only highlights the ability to guide prioritization of specific mutant alleles in important oncogenic pathways, but can also facilitate generation of novel hypotheses toward functional annotation and gene discovery.

## Discussion

REVEALER 2.0 provides a powerful approach to study mutational and copy number landscapes in a functional context. By using the *information* and *conditional information coefficient*, it strikes a good balance between weighting the features’ complementarity vs. their association with the functional target, and a good compromise between sensitivity and specificity. REVEALER 2.0 is particularly useful in cases where there is an accurate sample-per-sample functional profile representing a phenotype, activation profile, or biological state of interest, and a comprehensive characterization of genomic abnormalities in large datasets (n > 10,000). Functional annotation of individual alleles *via* traditional experimental approaches, either individually or systematically, can be time consuming and costly.^18,19^ Implementing REVEALER 2.0 can guide such characterization efforts and help prioritize the most promising alleles. The limitations of the original implementation of REVEALER, in terms of large execution times, limited its applicability to large datasets. However, the approach presented here, both the mathematical reformulation of the algorithm, and its efficient implementation, completely solves that problem and at the same time adds other capabilities and enables additional application modalities. This new implementation of REVEALER 2.0 addresses the unmet need to perform genome discovery efforts across datasets ever increasing in size and complexities, such as pan-cancer TCGA or those from the International Cancer Genome Consortium (ICGC), whole genome sequencing of somatic and germline data, as well as genomic data from single-cell profiling across hundreds of thousands of individual cells.

The application of REVEALER 2.0 to specific phenotypes associated with the activation of major oncogenes or oncogenic pathways (e.g. MAPK, β-catenin/WNT etc.) reveals a complex landscape of mutational alterations that has been hidden and is difficult to elucidate without the explicit functional analysis as provided by REVALER 2.0. Moreover, REVEALER 2.0 enables a high-resolution study of which specific mutant alleles behave as activating lesions either because of their high complement to other lesions present in the same or a related gene, or because of their high association with the functional phenotype.

REVEALER 2.0 “high-resolution” analysis of activating alleles and copy number alterations hold the promise of providing a much needed functional understanding of the mutational landscapes associated with major oncogenic pathways and processes. A better understanding of mutational landscapes in turn can provide critical benefits for precision oncology paradigms where, e.g., the sequencing profiles of tumor samples, can be interpreted at a higher level of resolution.

### Input file preprocessing for REVEALER 2.0 Analysis

REVEALER 2.0 has a companion file processing and preparation program that allows the user to generate customized input feature files from raw *maf* format files. In addition a user can generate input files by separating a specific feature into allele level, mutational function level, or matching features/alleles to targets and collapsing alleles that are highly correlated with the target. This function is highly customizable and flexible so that users can create input files tailored for specific analysis modalities for REVEALER 2.0.

### REVEALER 2.0 distribution

REVEALER 2.0 will be available as a standalone program and as a module in GenePattern (genepattern.org). Code and example notebooks can be obtained in https://github.com/yoshihiko1218/REVEALER2.

## Acknowledgments

This work was supported by grants: NIH U24 CA220341, NIH U24 CA248457, NIH R01 CA247551, NIH U01 CA217885, NIH R01 CA109467, NIH R01 CA226803, and a State of California’s *Initiative to Advance Precision Medicine CIAPM award* (OPR18112).

