## Supplemental Information for "Deciphering the Functional Roles of Individual Cancer Alleles Across Comprehensive Cancer Genomic Studies"

### Appendix

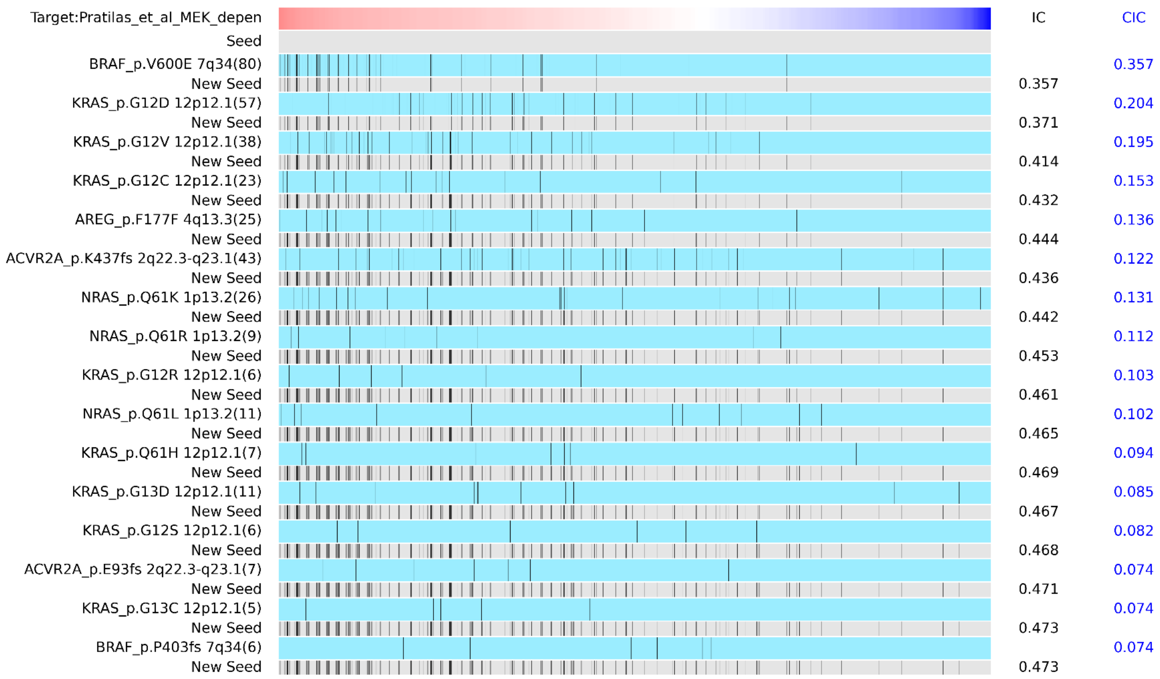


**Fig.S1. Individual alleles identified to match MEK inhibitor sensitivity profiles across CCLE dataset.**

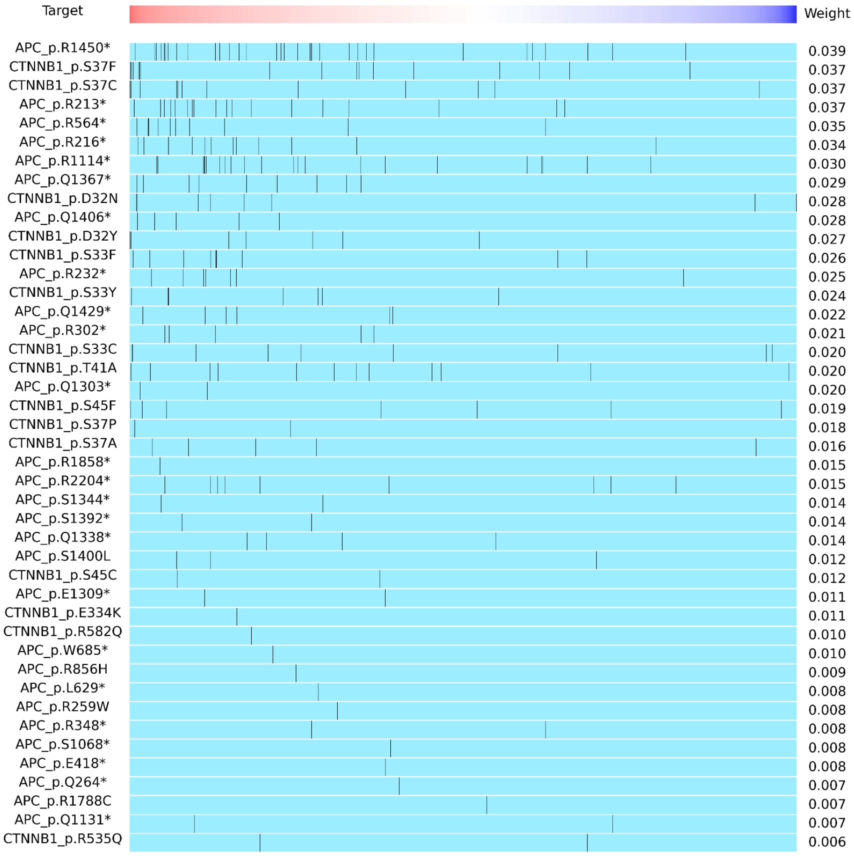

**Fig.S2. Distribution of shared allele between TCGA and CCLE for APC and CTNNB1 in TCGA.**

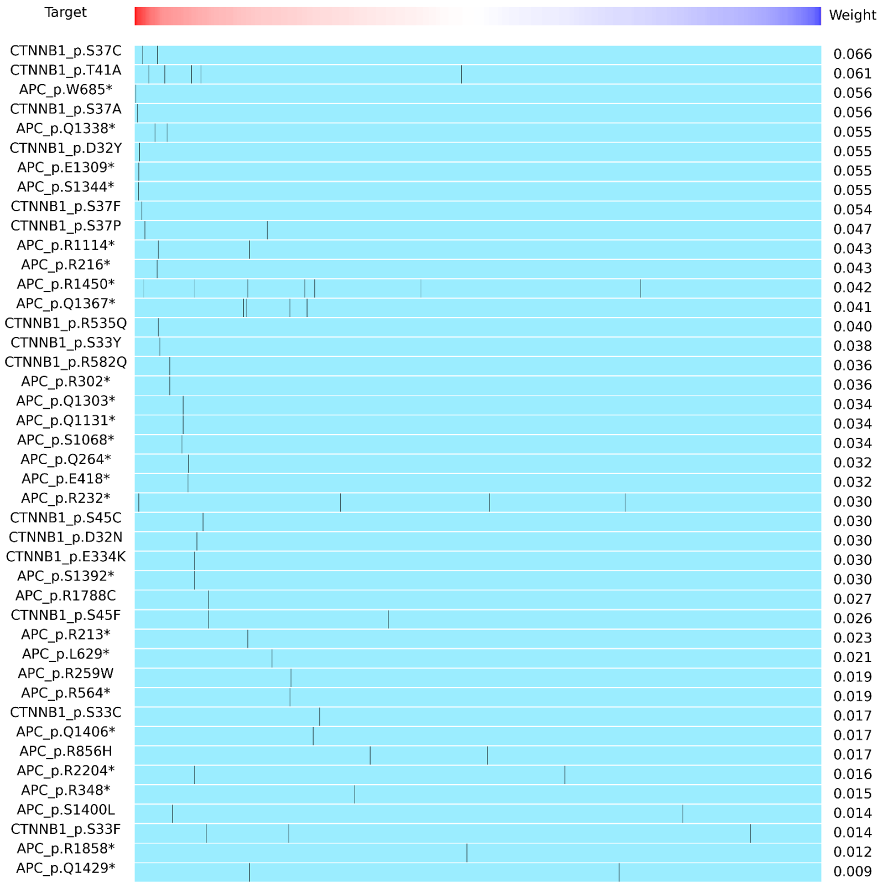


**Fig.S3. Distribution of shared allele between TCGA and CCLE for APC and CTNNB1 in CCLE.**

**Table S1. Selected shared allele between CCLE and TCGA for seed.**

| Allele | # in TCGA | # in CCLE | IC in TCGA | IC in CCLE |
|---|---|---|---|---|
| CTNNB1_p.S37P | 2 | 2 | 0.01773278059 | 0.04669580491 |
| CTNNB1_p.R535Q | 2 | 1 | 0.006313298942 | 0.04047482154 |
| APC_p.R1788C | 1 | 1 | 0.007280656828 | 0.0273283999 |
| APC_p.R213* | 18 | 1 | 0.03697079872 | 0.02300608758 |
| CTNNB1_p.R582Q | 1 | 1 | 0.01015598976 | 0.03620944376 |
| APC_p.R1858* | 1 | 1 | 0.01518934946 | 0.01216288835 |
| APC_p.R302* | 5 | 1 | 0.02067092654 | 0.03620944376 |
| APC_p.R1114* | 22 | 2 | 0.02986231983 | 0.04263805683 |
| APC_p.R564* | 10 | 1 | 0.0352552338 | 0.01888885031 |
| APC_p.E418* | 1 | 1 | 0.007585962942 | 0.03177155739 |
| APC_p.Q1303* | 2 | 1 | 0.01955126289 | 0.03401738705 |

| | | | | |
|---|---|---|---|---|
| APC_p.Q264* | 1 | 1 | 0.007412337262 | 0.03177290284 |
| APC_p.E1309* | 2 | 1 | 0.01136157653 | 0.0547640029 |
| CTNNB1_p.D32Y | 7 | 1 | 0.02662335567 | 0.05477540667 |
| APC_p.S1068* | 1 | 1 | 0.007586586107 | 0.03401346697 |
| CTNNB1_p.S45F | 7 | 2 | 0.019486764 | 0.02567861171 |
| APC_p.S1392* | 2 | 1 | 0.0140155695 | 0.02954586481 |
| APC_p.R856H | 1 | 2 | 0.008811332457 | 0.01688755204 |
| APC_p.R348* | 2 | 1 | 0.00801119406 | 0.0153380657 |
| APC_p.R232* | 8 | 4 | 0.02549773577 | 0.02971827908 |
| APC_p.S1344* | 2 | 1 | 0.01410892482 | 0.05476207206 |
| APC_p.L629* | 1 | 1 | 0.008434642013 | 0.02091499396 |
| APC_p.Q1429* | 6 | 2 | 0.02176193628 | 0.009005785645 |
| APC_p.R2204* | 9 | 2 | 0.0147177421 | 0.01574616193 |
| APC_p.R1450* | 33 | 7 | 0.03871992573 | 0.04225225117 |
| CTNNB1_p.T41A | 12 | 5 | 0.02036068586 | 0.06132758776 |
| APC_p.S1400L | 3 | 2 | 0.01198226688 | 0.01430999111 |
| CTNNB1_p.S45C | 2 | 1 | 0.01182204581 | 0.0295709936 |
| CTNNB1_p.S37F | 17 | 1 | 0.03722970814 | 0.05386526 |
| APC_p.W685* | 1 | 1 | 0.009677875381 | 0.05647521057 |
| APC_p.Q1338* | 4 | 2 | 0.0140093339 | 0.05495506511 |
| APC_p.Q1406* | 5 | 1 | 0.02773061456 | 0.01700564404 |
| APC_p.Q1131* | 2 | 1 | 0.006799324145 | 0.03401738705 |
| CTNNB1_p.S33Y | 7 | 1 | 0.02392018653 | 0.03837452735 |
| CTNNB1_p.S37C | 14 | 2 | 0.03706430059 | 0.0658468972 |
| APC_p.R259W | 1 | 1 | 0.008100922344 | 0.01888909233 |
| CTNNB1_p.E334K | 1 | 1 | 0.01066077036 | 0.02954586481 |
| CTNNB1_p.S33C | 9 | 1 | 0.02039605843 | 0.01701396992 |
| APC_p.Q1367* | 9 | 4 | 0.0291661876 | 0.04101448313 |
| APC_p.R216* | 13 | 1 | 0.03413403356 | 0.04256983572 |
| CTNNB1_p.S33F | 9 | 3 | 0.02631809628 | 0.01351061918 |
| CTNNB1_p.D32N | 8 | 1 | 0.02807640612 | 0.02955497267 |
| CTNNB1_p.S37A | 5 | 1 | 0.01631909267 | 0.05558252128 |
